# Matrix stiffness regulates Notch signaling activity in endothelial cells

**DOI:** 10.1101/2022.04.20.488899

**Authors:** Maibritt Kretschmer, Rose Mamistvalov, David Sprinzak, Angelika M. Vollmar, Stefan Zahler

**Affiliations:** Department of Pharmacy, Pharmaceutical Biology, Ludwig-Maximilians-Universität München, Butenandtstraße 5-13, 81377 Munich, Germany; The George S. Wise Faculty of Life Sciences, Tel Aviv University, Tel Aviv 69978, Israel

**Keywords:** substrate stiffness, mechanosensing, endothelial cells, Notch1, Dll4

## Abstract

The Notch signaling pathway plays a critical role in many developmental and disease related processes. It is widely accepted that Notch has a mechano-transduction module that regulates cleavage of the receptor. However, the role of biomechanical properties of the cellular environment on this module and on Notch signaling in general is still poorly understood. During angiogenesis, differentiation into tip and stalk cells is regulated by Notch. The endothelial cells in this process respond to biochemical and mechanical cues triggered by local stiffening of the ECM. Here, we investigated the influence of substrate stiffness on the Notch signaling pathway in endothelial cells. Using stiffness tuned PDMS substrates we show that Notch signaling pathway activity inversely correlates with the physiologically relevant substrate stiffness, with increased Notch activity on softer substrates. We show that trans-endocytosis of the Notch extracellular domain, but not the overall endocytosis, is regulated by substrate stiffness. Furthermore, we could show that integrin cell-matrix connections are both stiffness-dependent and influenced by Notch. Cadherin mediated cell-cell adhesion and Notch, however, influence each other in that basal Notch signaling is cell-cell contact dependent, but inhibition of the Notch signaling pathway also results in a reduction of VE-cadherin levels. We conclude that mechano-transduction of Notch activation depends on substrate stiffness highlighting the role of substrate rigidity as a modulator of Notch signaling. This may have important implications in pathological situations, such as tumor growth, associated with stiffening of the extracellular matrix.

## Introduction

The Notch signaling pathway plays a crucial role in many tissues, regulating cell fate decisions, cell cycle progression and apoptosis during tissue development and maintenance [1, 2]. Notch signaling is induced by a Notch ligand (Delta-like 1, 3 or 4, Jagged 1 or 2) interacting with one of four types of Notch receptors (Notch1-4) of an adjacent cell (trans interaction), which triggers a signaling cascade in the receiver cell [2–4]. Receptor-ligand binding leads to two proteolytic cleavage events, resulting first in separation of the Notch extracellular domain (NECD), and subsequently in release of the Notch intracellular domain (NICD) [5–7]. The NECD is pulled into the ligand-presenting cell (sender cell) along with the ligand in the course of trans-endocytosis, while the NICD is translocated to the nucleus of the receiver cell, where it associates with the transcriptional regulator RBPJ, activating the Notch target genes such as Hairy-Enhancer of Split (HES) and HES-related proteins [1, 7, 8].

During Notch activation, mechanical forces generated by endocytosis in the sender cell are required for opening the negative regulatory region (NRR) in the Notch receptor, leading to the first proteolytic cleavage event [8, 9]. However, the influence of mechanical cues from the surrounding matrix on this process (i.e. mechanosensitivity) is poorly studied.

The Notch signaling pathway is instrumental for angiogenesis, regulating the selection of tip and stalk cells [10, 11]. Of the four Notch homologs and five Notch ligands that have been identified in mammals the Notch receptor Notch1 and the ligands Dll4 and Jag1 are mainly involved in tip/stalk cell specification [2, 11, 12]. Tip cells are characterized by increased Dll4 expression mediated by VEGF-VEGFR2 signaling [11, 12]. Stalk cells, on the other hand, exhibit increased Notch1 expression, associated with the high Dll4 level in the tip cells. Activation of the Notch signaling pathway in stalk cells by Dll4 expression in tip cells, induces cell proliferation of stalk cells, resulting in extension of sprouts and lumen formation [12, 13]. While the biochemical cues during angiogenesis are well studied, the mechanical forces during angiogenesis are only beginning to be understood [14, 15]. Besides the shear force due to the blood flow, most of the mechanical cues during angiogenesis are exerted by the ECM, caused by matrix structural changes that can locally alter the ECM stiffness and density [16–18]. However, little is known about the mechanical influences via the matrix on the signaling pathways involved in angiogenesis, like the Notch signaling pathway.

In this study, we investigate the influence of different physiologically relevant substrate stiffnesses on the Notch signaling pathway and related pathways in endothelial cells such as YAP and adhesion dependent signaling. Using different stiffness-tuned silicone substrates and two different modes of Notch activation with DLL4 as a ligand (via cell-cell contact or a protein coat), we show that activation of the Notch signaling pathway is stiffness dependent. Furthermore, we investigate the influence of cell-cell contacts on Notch activation and the mutual influence of Notch and cell-cell or cell-matrix adhesion proteins, as well as the cross-talk between Notch and YAP signaling, a well-known mechanosensitive pathway [2]. Together, our results demonstrate a mechanosensitivity of the Notch signaling pathway, likely associated with the process of trans-endocytosis, suggesting a second mechanical aspect of the Notch signaling pathway besides the pulling force generated by the ligand presenting cell.

## Material and Methods

### Cell culture and co-cultures

Human Umbilical Vein Endothelial Cells (HUVECs) were obtained from Promocell (Heidelberg, Germany), Human Microvascular Endothelial Cells (HMECs) were a gift from Dr. Guido Jürgenliemk (University Regensburg, Germany). Cells were cultured in ECGM medium (PELO Biotech, Planegg, Germany) including the supplement kit, 10% heat inactivated fetal calf serum (FCS, PAA Laboratories GmbH, Pasching, Austria), 250 μg/ml amphotericin B and 10,000 U/ml penicillin/streptomycin (PAN Biotech, Aidenbach, Germany) at 37°C and 5% CO_2_ under constant humidity. Cells were maintained until passage 6 for the respective experiments.

Mouse Cardiac Endothelial Cells (MCECs) were a gift from Andreas Fischer (DKFZ, Heidelberg, Germany) and maintained in DMEM medium supplemented with 10% FCS (PAA Laboratories GmbH, Pasching, Austria) and 200mM L-glutamine (Thermo Fisher Scientific, Waltham, MA). Two MCEC cell lines were used: a wild type cell line (MCEC-WT) as well as a cell line overexpressing Dll4 coupled to a mCherry reporter (MCEC-Dll4-mCherry). At confluence, cells were split and cultivated with continuous passage.

All applied surfaces and substrates were coated with collagen G (10μg/ml in PBS, MATRIX BioScience GmbH, Mörlenbach, Germany), ensuring uniform adhesion.

HUVEC/MCEC co-cultures were suspended in ECGM medium; MCEC/MCEC co-cultures in DMEM medium.

### Generation of stable cell lines

The generation of MCEC TetR-TO-Dll4-mCherry (MCEC-Dll4-mCherry) was done in two steps. First, MCEC-TetR were generated by stable transfection of the pcDNA6-TR-Blast plasmid (gifted from the Bjorkman lab) and selection of single clones using 10ug/ml Blasticidine (H-1012-PBS; A.G. Scientific Inc., San Diego, CA). Then, MCEC-TetR were transfected with the pcDNA5-TO-hDll4-mCherry plasmid and were placed under another selection with 250ug/ml Hygromycin (B-1247-SOL; A.G. Scientific Inc., San Diego, CA). Single colonies were picked and tested for fluorescence under conditions of with and without 100ug/ml Doxycycline (Sigma-Aldrich, St. Louis, MO).

### Polydimethylsiloxane (PDMS) gels

PDMS substrates were prepared using the Sylgard™ 184 silicone elastomer kit (Dow Corning, Midland, MI). PDMS base was mixed with a ratio of 10%, 2% or 1.3% curing agent and was transferred to 8-well or 2-well μ-slides from ibidi (Martinsried, Germany). Gels were degassed in a desiccator for 10-15 min and polymerized in a compartment drier at 60°C for 20h, resulting in substrate stiffnesses of 70kPa, 1.5kPa and 0.5kPa. PDMS substrates were hydrophilized with oxygen plasma at 0.3 mbar for 3 min using the Zepto plasma system (Diener electronic GmbH & Co. KG, Ebhausen, Germany). Stiffnesses were determined after hydrophilization by rheometric measurements with the Modular Compact Rheometer MCR 100 (Physica, Stuttgart, Germany).

### Plasmids and transfections

The TP1-Luc construct was first published by Minoguchi et al [19]. pRL-SV40P Renilla was a gift from Ron Prywes (Addgene plasmid # 27163; https://www.addgene.org/27163/. The pcDNA3-hN1-citrine was constructed from pcdna3-hNECD-G4esn-cit described in previous work (Sprinzak et al., 2010, [3]). The G4esn was replaced with the full length of human Notch1. pcDNA5-TO-hDll4-mCherry was described by Sprinzak et al. mCitrine-VE-Cadherin-N-10 was a gift from Michael Davidson (Addgene plasmid # 56319; http://n2t.net/addgene:56319; RRID:Addgene_56319).

HUVECs were transfected transiently using either the Targefect-HUVEC™ transfection kit (Targeting Systems, El Cajon, CA) or the Cell Line Nucleofector™ Kit V with the program A-034 (Lonza Group AG, Basel, Switzerland) according to manufacturer’s protocol.

MCECs were transiently transfected with the FuGENE® Transfection Regent (Promega, Madison, WI) following manufacturer’s instructions.

Transfected cells were incubated for 24h before further assays or evaluations were applied.

### Reporter gene assay

Notch-responsive luciferase reporter assays were performed 24h subsequent to co-transfection of endothelial cells with the CSL-binding plasmid TP1-Luc and Renilla. Firefly and Renilla vector levels were applied in a ratio 10:1. Using the Dual-Luciferase® Reporter Assay System (Promega, Madison, WI) and the Orion II microplate luminometer equipped with Simplicity analysis software (Berthold Technologies, Bad Wildbad, Germany) luciferase levels were determined. Firefly RLUs were normalized to the Renilla control.

### Trans-endocytosis assay

Cells grown to 80% confluency (HUVECs or MCEC-WTs) were transiently transfected separately with the plasmids pcDNA3-hN1-citrine and pcDNA5-TO-hDll4-mCherry and incubated on plastic for 24h at 37°C under 5% CO_2_. Cells were then washed, detached, and reseeded together in a co-culture ratio 1:1. Doxycycline was added during reseeding in a concentration of 100 ng/ml for activation of the Dll4-mCherry expression. After 6h incubation at 37°C under 5% CO_2_, co-cultures were fixed with 4% methanol free formaldehyde (Thermo Fisher Scientific, Waltham, MA) in PBS for 10 min. Samples were washed twice with PBS and sealed with FluorSave mounting medium. Trans-endocytosis was visualized by confocal microscopy. Quantification was performed by analysis of co-localized areas and its intensity using ImageJ.

### Transferrin endocytosis assay

Cells grown to 100% confluence were washed once with PBS including Ca^2+^ and Mg^2+^ (PBS^+^) and incubated for 10 min with 5μg/ml Transferrin, Alexa Flour 488 conjugate (Life Technologies, Carlsbad, CA) at 37°C under 5% CO_2_. Cells were washed once with room temperature acid wash medium and then fixed with 4% methanol free formaldehyde (Thermo Fisher Scientific, Waltham, MA) in PBS for 10 min. Samples were washed twice with PBS and sealed with FluorSave mounting medium. Transferrin uptake was visualized by confocal microscopy.

### Antibodies, compounds, and staining reagents

The following primary antibodies used in this study were directed against: Cleaved Notch1 (Val1744) rabbit mAB IgG, 4147 (Cell Signaling Technology, Danvers, MA); Integrin β1 rabbit pAB, 4706 (Cell Signaling Technology, Danvers, MA); Anti-Integrin beta 1 (12G10) mouse mAB IgG1, ab30394 (Abcam, Cambridge UK); YAP (D8H1X) XP® rabbit mAB IgG, 14074 (Cell Signaling Technology, Danvers, MA); VE-Cadherin rabbit pAB, 2158 (Cell Signaling Technology, Danvers, MA). Following secondary antibodies were applied in this study: Alexa Fluor 488-conjugated goat anti-mouse IgG (H+L), A-11001; Alexa Fluor 488-conjugated goat anti-rabbit IgG (H+L), A-11008; Alexa Fluor 647-conjugated chicken anti-rabbit IgG (H+L), A-21443 (all from Thermo Fisher Scientific, Waltham, MA).

DAPT was purchased from Sigma Aldrich, dissolved in DMSO to a stock concentration of 10mM and used at working concentrations of 25μM. DMSO controls were performed with the appropriate DMSO concentrations. EGTA was obtained from Sigma Aldrich and applied at 5mM. The VE-cadherin blocking antibody clone BV6 was obtained from Merck Millipore, Darmstadt, Germany and was diluted 1:10 for use (50μg/ml). Hoechst 33342 was purchased from Sigma Aldrich, solved in PBS, and applied at a final concentration of 10μg/ml. FluorSave™ Reagent was purchased from Merck Millipore (Darmstadt, Germany).

### Immunofluorescence Staining

Cells were washed once with phosphate buffered saline containing Ca^2+^ and Mg^2+^ (PBS^+^) and were fixed with 4% methanol free formaldehyde (Thermo Fisher Scientific, Waltham, MA) in PBS for 10 min. Fixation was followed by a brief washing with PBS and cell permeabilization with 0.1% Triton X-100 in PBS for 10 min. After another brief washing with PBS, nonspecific binding sites were blocked with 5% BSA (Roth, Karlsruhe, Germany) in PBS for 60 min. Cells were then incubated with the primary antibody diluted in PBS with 1% BSA (1:200) overnight at 4°C. Next, samples were washed 3×10 min with 1% BSA in PBS, then incubated with the secondary antibody (1:400) and Hoechst 33342 (1:100) for nuclear counter stain, again diluted in PBS with 1% BSA for 1h at room temperature. Cells were washed again 2×10min with 1% BSA in PBS and once 10 min with PBS and were sealed with FluorSave mounting medium.

### Laser scanning confocal microscopy

Confocal images were taken with a Leica TCS SP8 microscope equipped with an HC PL APO CS2 63x/1.4 oil objective and photomultiplier (PMT) or HyD detectors, using the LAS X core software. In sequential scanning mode two frames were acquired for every channel with a scanning speed of 400 Hz and the pinhole size set to 1.0 airy units. Following excitation laser lines were applied: 405 nm, 488 nm, and 647 nm.

### Fluorescence recovery after photobleaching (FRAP)

HUVECs were transiently transfected with mCitrine-VE-Cadherin-N-10, seeded directly on different substrates (plastic and PDMS) and incubated for 24h at 37°C under 5% CO_2_. The FRAP assay was conducted with the Leica TCS SP8 SMD microscope with the HC PL APO CS2 63x/1.4 NA oil objective and the heating and gas incubation system from OCO Lab (Naples, Italy) ensuring constant 37°C under 5% CO_2_ and 80% humidity. Using the LAS X Core Software, the FRAP settings were adjusted to one pre-bleach iteration, 20 bleach iterations, five post-bleach iterations with 30 sec intervals and seven with 60 sec intervals. Images were taken with a pinhole size adjusted to 1.0 airy units and a scanning speed of 400 Hz. The line 488 (argon) and the PMT detector were applied.

### Data analysis and statistics

All confocal images were analyzed using ImageJ version 1.53c. Pearson’s R coefficients were determined using the Coloc2 plugin for ImageJ. Intensity ratios nuclear/cytoplasmatic were evaluated with the Intensity Ratio Nuclei Cytoplasm Tool plugin for ImageJ. Expression patterns were determined by segmentation and skeletonizing of the images.

All data was derived from three independent experiments represented as the mean ± SEM. Statistical analysis (ordinary one-way ANOVA with Dunnett’s multiple comparison test, two-way ANOVA with Tukey’s multiple comparison test) were processed using GraphPad Prism 9.2.0

## Results

### Decreased substrate stiffness increases Notch signaling activity in endothelial cells

To investigate the mechanosensitivity of the Notch signaling pathway, synthetic substrates with defined stiffness in a range from 0.5 to 70 kPa were applied. Endothelial cells (HUVEC and MCEC-WT cells) were used as prototypic models for Notch signaling. The Notch signaling pathway was activated either via cell seeding onto an rhDll4 coat or by co-culture of HUVEC cells (receivers) with Dll4 overexpressing MCECs (MCEC-Dll4-mCherry, senders). Due to the dependence of the primary endothelial cells in in vitro culture on a protein coat, all surfaces were coated with collagen G unless indicated otherwise. Analysis of a Notch reporter gene assay showed a continuous increase in Notch transcriptional activity on softer substrates both after activation by cell-cell contact using the Dll4 overexpressing cell line, and after activation by a rhDll4 protein coating. (Figure 1A and 1C). These findings were supported by intensity analysis of the nuclear localization of the Notch intracellular domain (NICD) after immunofluorescence staining: the softer the substrate, the higher the NICD intensity in the Notch receiver cells upon stimulation (Figure 1B and 1D). Thus, Notch activation, both by plate bound Dll4 and by co-cultured sender cells exhibits mechanosensitivity. Activation of the Notch signaling pathway by co-culture with MCEC-Dll4s in different ratios using HUVEC cells and MCEC-WTs as signal receiver cells showed a 1:1 ratio to be optimal. With a higher amount of sender cells overall signal intensity decreased (Supplementary Figure S1A and B). Human Microvascular Endothelial Cells (HMEC) were used as an additional endothelial cell model. After co-culture with MCEC-Dll4-mCherrys they showed results comparable to HUVECs and MCECs, and the same mechanosensitivity (Supplementary Figure S1C). The efficiency and reproducibility of coating of the PDMS substrates with rhDll4 were controlled in an availability assay by immunostaining, which showed that the rhDll4 coat was evenly distributed, and showed a comparable intensity on all substrates, (Supplementary Figure S1D).

**Figure 1.**
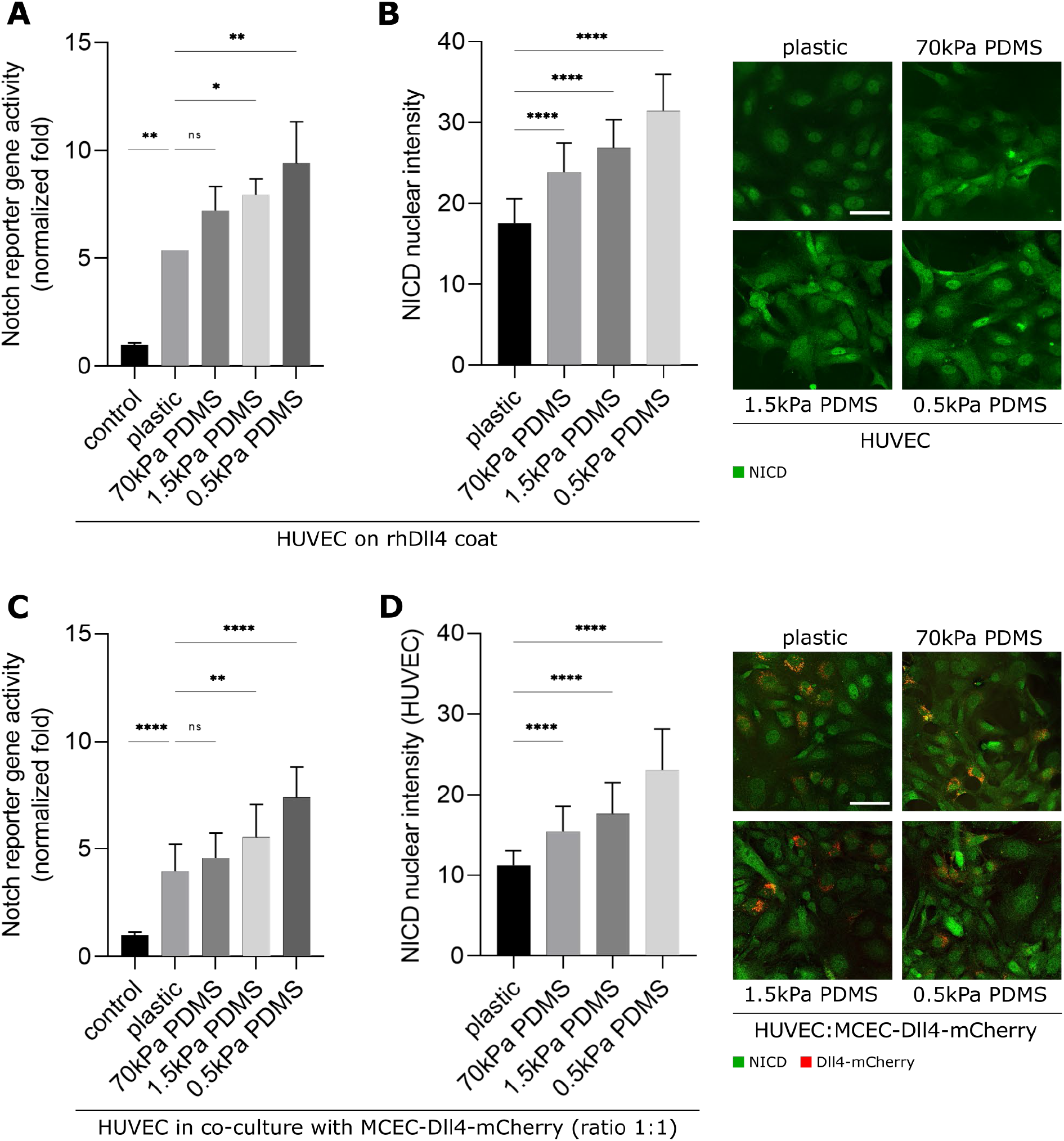
Notch activity in endothelial cells increases on softer substrates. (A,B) Induction of Notch activity by coating with rhDll4. (A) Normalized fold Notch activity in confluent HUVEC cells on PDMS substrates with different stiffnesses, determined by detection of cellular luciferase levels under control of the TP1-luc Notch reporter. (B) Nuclear NICD intensities in HUVEC cells seeded onto different plastic and PDMS substrates. Left panel: quantitative evaluation of nuclear fluorescence intensity of NICD; right panel: representative images of cells stained for NICD (shown in green, scale bar 50μm). Mean values ± SEM, *P<0.1, **P<0.01, ****P<0.0001 (one way ANOVA followed by Dunnett’s multiple comparison test). (C,D) Induction of Notch activity by co-culture with MCEC-Dll4-mCherry cells on a collagen G coat. (C) Normalized fold Notch activity in confluent HUVEC/MCEC-Dll4-mCherry co-cultures on substrates with different stiffnesses, determined by detection of cellular luciferase levels under control of the TP1-luc Notch reporter. (D) Left panel: nuclear NICD intensities in HUVEC cells seeded in co-culture with MCEC-Dll4-mCherry cells on different plastic and PDMS substrates. Intensities are shown in a bar graph on the left panel, quantified in ≥300 single cells derived from 3 independent experiments; right panel: representative images of cells stained for NICD (shown in green, mCherry-Dll4 reporter of MCEC-Dll4-mCherry cells shown in red, scale bar 50μm). Mean values ± SEM, **P<0.01, ****P<0.0001 (Tukey’s corrected one way ANOVA).

### Active integrin β1 increases on softer substrates and is influenced by Notch

Since interaction with the ECM is known to be mediated by integrin signaling, we next wanted to check if Notch signaling affects or is affected by integrins [7]. Integrin β1 represents the largest subchain of integrins and is involved in several biological processes such as adhesion, migration and cell cycle regulation [20]. Due to the involvement of β1 subchains in cell-ECM interaction, integrin β1 plays a major role especially in ECs [21].To investigate, whether mechanosensitivity of Notch lies up- or downstream of integrin signaling, we quantified total and activated β1 integrin levels with and without pretreatment of cells with the Notch inhibitor DAPT on the different substrates. Previous studies have already shown that activated but not overall integrin levels are substrate-dependent and increased on softer substrates [22]. Accordingly, in our model overall intensity of integrin β1 in HUVEC cells does not change on the different substrates (Figure 2A). In contrast, the softer the substrate, the more integrin β1 is activated (Figure 2B). Upon blocking of basal Notch1 cleavage, and, thus, downstream of Notch signaling, integrin β1 activation decreases substantially, although the correlation between softer substrates and increased integrin β1 activation remains to a small degree (Figure 2C). Thus, Notch signaling occurs upstream of integrin activation, as far as mechanosignaling is concerned. This result was confirmed in the second endothelial cell line MCEC-WT as well (Supplementary Figure S3A).

**Figure 2.**
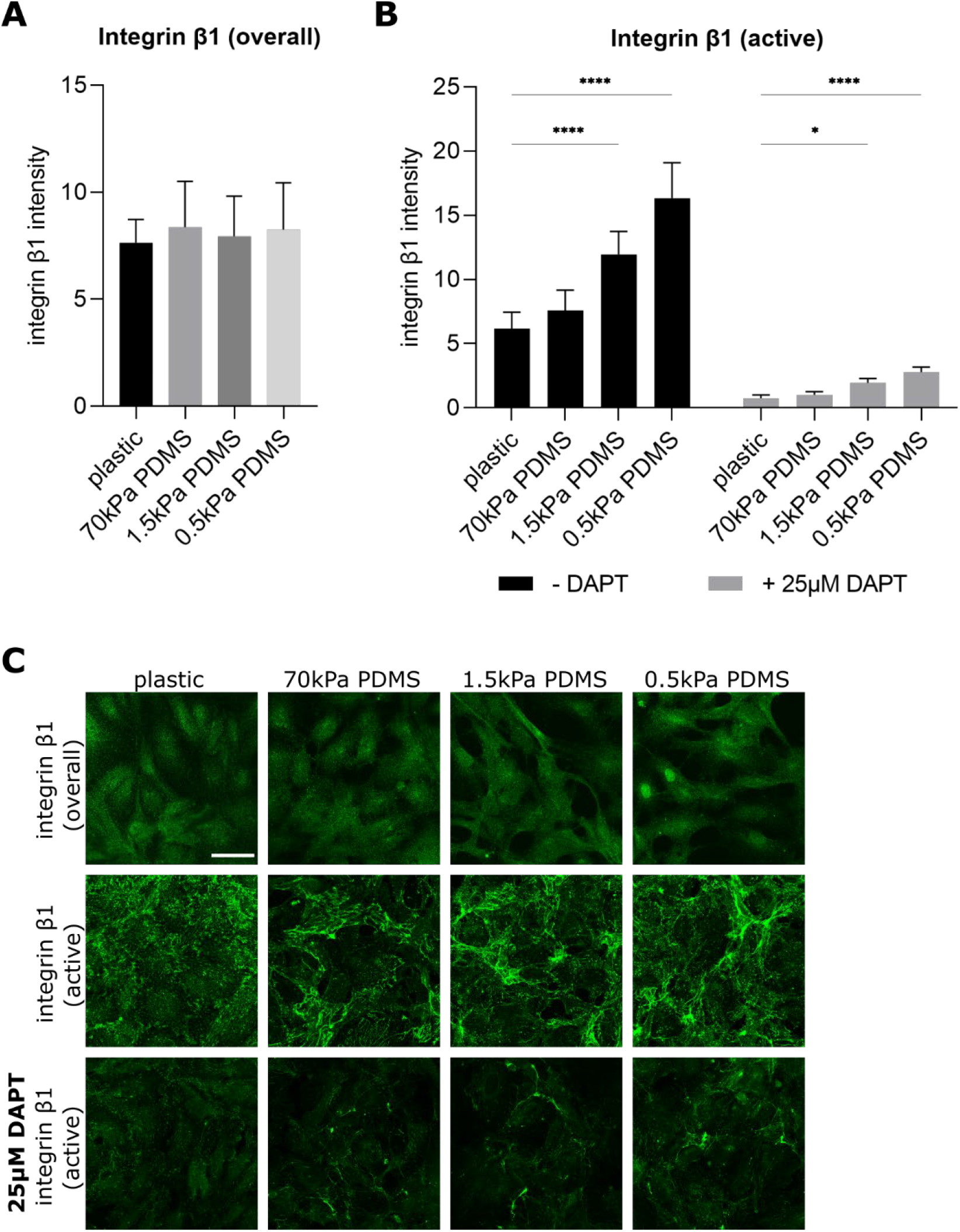
Integrin β1 activity relates to substrate stiffness and is influenced by Notch. (A) Integrin β1 intensities in HUVECs. Cells were seeded on varying substrate stiffness and were stained for total integrin β1. Mean intensities ± SEM are depicted in bar graphs (one-way ANOVA followed by Tukey’s multiple comparison test). (B) Integrin β1 (active) intensity in HUVECs. Cells were seeded on varying substrate stiffness, treated with 25μM DAPT for 24h and stained for the activated form of integrin β1. Data is presented in a bar plot and compared with the integrin β1 intensities without DAPT treatment (mean intensity ± SEM, Tukey’s corrected two-way ANOVA, *P<0.1, ****P<0.0001). (C) Representative images of HUVEC cells on plastic and PDMS substrates +/- DAPT treatment stained for total or activated integrin β1 (green) are shown (scale bar 50μm).

### Yes-associated protein (YAP) signaling and Notch are inversely mechano-regulated

The role of the Yes-associated protein YAP as a mechano-transducer and the related mechano-sensitivity of the YAP/TAZ signaling pathway are well known [23, 24]. We and others have shown that YAP activity decreases on soft substrates [25]. To asses a possible YAP/Notch crosstalk in endothelial cells, we performed immunofluorescence staining in HUVEC cells seeded on plastic and PDMS substrates, again with and without addition of DAPT (25μM, 24h). As expected, our results show progressively decreased nuclear YAP intensity in HUVEC cells on softer substrates, demonstrating the mechano-sensitivity of the YAP/TAZ signaling pathway in our cells (Figure 3A). However, inhibition of the Notch signaling pathway has no clear effect on nuclear YAP intensity, only on the softest substrate of 0.5kPa Notch inhibition rescued YAP activation to some degree (Figure 3B). Consistent results were obtained with the control cell line MCEC-WT, shown in Supplementary Figure S3B. Since YAP and Notch activity are inversely regulated by substrate stiffness, there seems to be no direct crosstalk between these two signaling pathways.

**Figure 3.**
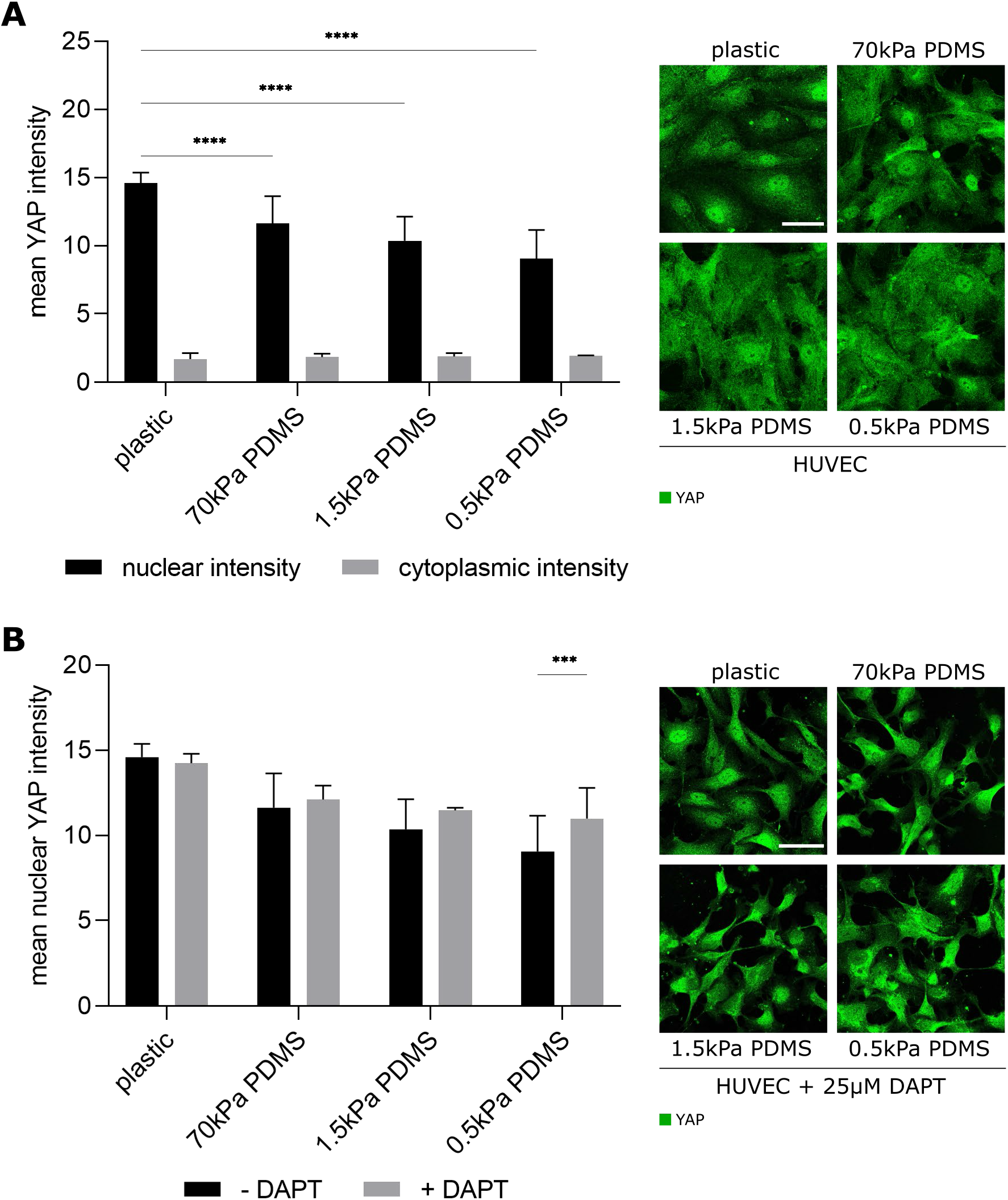
Nuclear YAP intensity is reduced on softer substrates but is only marginally affected by Notch inhibition. (A) Nuclear and cytoplasmic YAP intensities. HUVEC cells were seeded on varying substrate stiffness and were stained for YAP. Intensities were analyzed with the Intensity Ratio Nuclei Cytoplasm Tool plugin for ImageJ and are presented in a bar graph on the left panel (mean ± SEM, Sidak’s corrected two-way ANOVA, ****P<0.0001). Representative images of immunofluorescence staining are shown on the right panel, with YAP in green (scale bar 50μm). (B) Nuclear YAP intensities with and without Notch inhibition. HUVEC cells were seeded on varying substrate stiffness, treated with 25μM DAPT for 24h and stained for YAP. Intensities ± SEM of untreated and treated cells are summarized in a bar graph on the left panel (two-way ANOVA followed by Sidak’s multiple comparison test, ***P<0.001). Representative images of immunofluorescent stained HUVEC cells treated with DAPT are shown on the right panel with YAP in green (scale bar 50μm).

### VE-cadherin levels and trafficking to cell-cell borders are not affected by substrate stiffness, although the morphology of cell-cell contacts changes

Since the Notch signaling pathway is a contact-dependent pathway, we examined the influence of substrate stiffness on the major endothelial adhesion molecule VE-cadherin. No correlation between VE-cadherin intensity and substrate stiffness was detected. The junction patterns, however, showed stiffness-related changes. The softer the substrate, the less typically branched and interlinked junction pattern are observed at the cell-cell contacts, and a continuous VE-cadherin junction with larger intensity area without branches or comb-like structures is visible instead, as shown in the representative images (Figure 4A and 4B). We investigated the influence of substrate stiffness on VE-cadherin trafficking at cell-cell contacts using a FRAP assay. HUVEC cells were transiently transfected with a citrine-coupled VE-cadherin plasmid and seeded on PDMS substrate with different stiffness. VE-cadherin recovered to the same extent and with the same kinetics on all substrates and, therefore, no significant differences in the recovery half time were observed (Figure 4C). Thus, the altered junction patterns on the different substrates does not depend on, or affect VE-cadherin kinetics.

**Figure 4.**
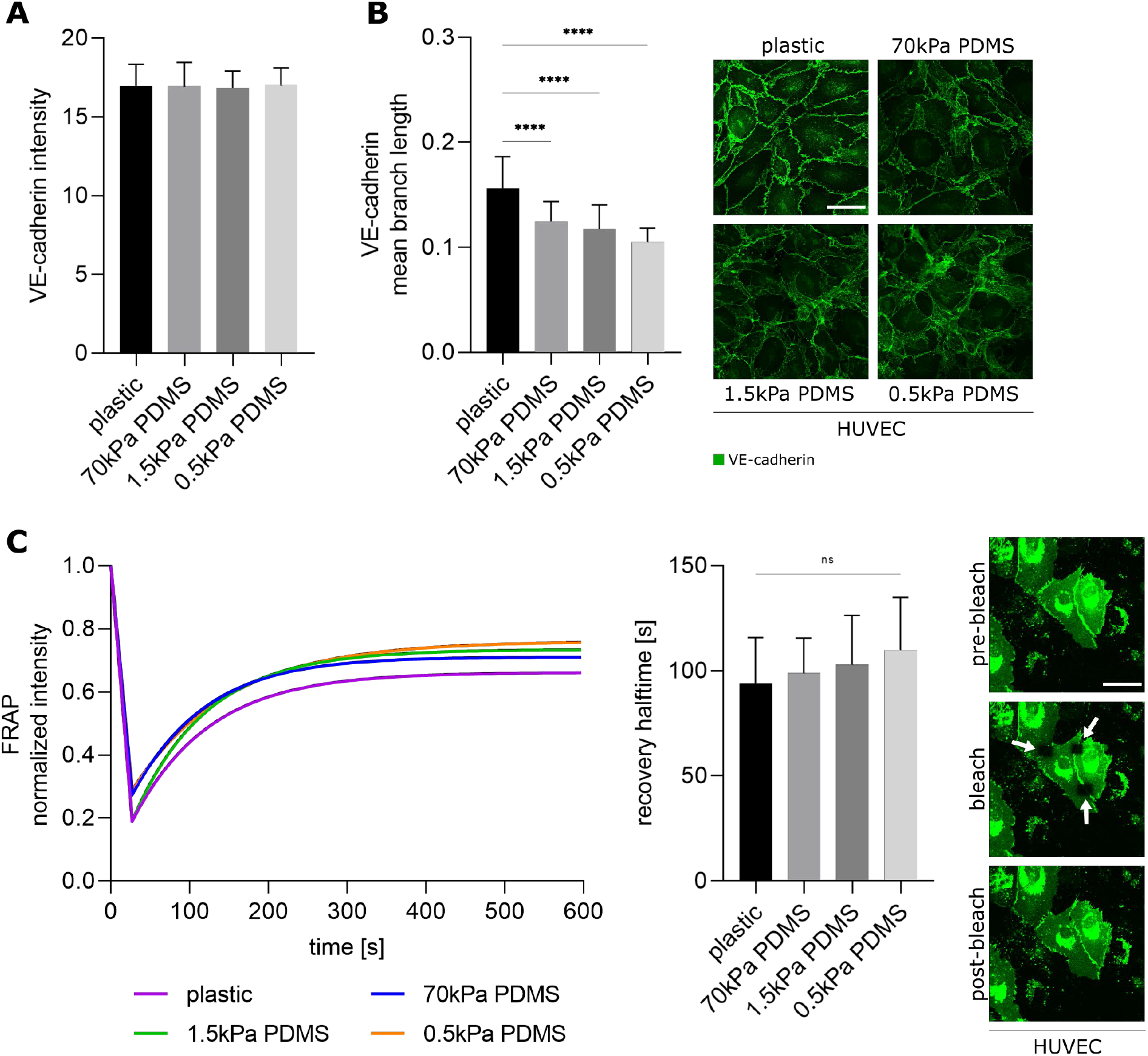
Softer substrates change VE-cadherin junction morphology but not VE-cadherin levels or its mobility. (A) VE-cadherin intensity in HUVEC cells. (B) VE-cadherin junction analysis in HUVECs, quantified by evaluation of the mean branch length. Cells were seeded on varying substrate stiffness and were stained for VE-cadherin. Intensity and junction analysis are presented in bar graphs (mean ± SEM, one-way ANOVA followed by Tukey’s multiple comparison test, ****P<0.0001). Representative images of immunofluorescence staining are shown on the right panel, with VE-cadherin in green (scale bar 50μm). (C) Analysis of VE-cadherin mobility at the cell-cell border in HUVEC via a FRAP assay. Cells were transiently transfected with mCitrine-VE-Cadherin-N-10 and seeded on varying substrate stiffness. FRAP was conducted with the Leica photo bleaching module. VE-cadherin recovery is plotted over time (left panel) and quantified as recovery half time (middle panel), shown in a bar graph (mean ± SEM, Tukey’s corrected one-way ANOVA). Representative images of the three FRAP steps, with citrine-coupled VE-cadherin in green are shown on the right panel (scale bar 50μm).

### Endogenous Notch activity is affected by breaking up of cell-cell contacts

To investigate the importance of existing cell-cell contacts for basal Notch activity, we seeded HUVEC cells on substrates of different stiffness and destabilized VE-cadherin adhesion molecules first acutely by EGTA and then inhibited them in a prolonged manner using a VE-cadherin blocking antibody. Treated cells were stained for NICD and its intensity was analyzed in the nuclei as readout for Notch activity. Since studies show that EGTA can affect the structural activity of Notch1, leading to receptor cleavage and activation [26, 27], we checked the effect of EGTA on NICD levels in our model first. Destabilization of cell-cell contacts leads to a reduction in nuclear NICD levels. Further blocking of VE-cadherin enhances this effect (Figure 5A) independently of substrate stiffness. The representative images (0.5kPa PDMS) confirm the quantitative analyses and show that the treatment changes the cell morphology and causes the cells to drift apart, leading to diminished cell-cell contacts. Furthermore, we also investigated the effect of Notch on VE-cadherin by treatment with DAPT (25μM, 24h). The results show a significant reduction in VE-cadherin intensity after Notch inhibition on all substrates. The junction patterns, however, do not change with the addition of DAPT (Figure 5B). Thus, changes in VE-cadherin morphology due to substrate stiffness seem to be independent of Notch signaling, while overall VE-cadherin expression is not.

**Figure 5.**
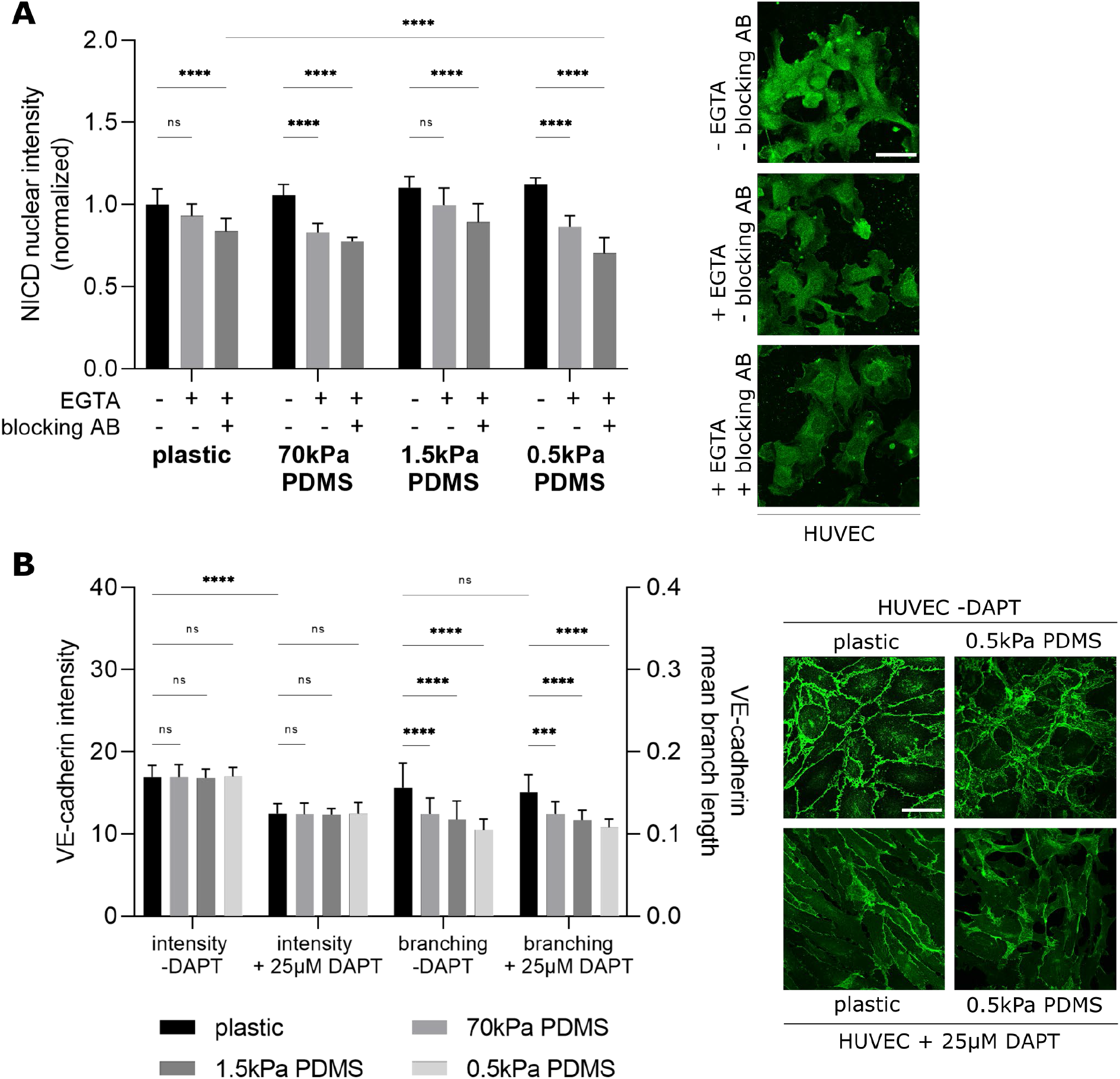
Notch and VE-cadherin influence each other: basal Notch activity is reduced by VE-cadherin blocking and VE-cadherin intensity is decreased by Notch inhibition. (A) Nuclear NICD intensity in HUVEC after cell-cell contact inhibition. Cells were seeded on substrates of varying stiffness without activation of the Notch signaling pathway, treated with EGTA and VE-cadherin blocking antibody and stained for NICD. Intensities are compared in a bar graph (mean ± SEM, Tukey’s corrected two-way ANOVA, ****P<0.0001) on the left panel. Representative images of HUVEC cells on 0.5kPa PDMS after immunofluorescence staining are shown on the right panel, with NICD in green (scale bar 50μm). (B) VE-cadherin intensity and junction analysis in HUVEC with and without Notch inhibition. Cells were seeded on varying substrate stiffness, treated with 25μM DAPT for 24h and stained for VE-cadherin. Bar plots were generated by evaluation of intensity and mean branch length, displayed on the left panel (mean ± SEM, one-way ANOVA followed by Tukey’s multiple comparison test, *P<0.1, **P<0.01, ****P<0.0001). Representative images of HUVECs on plastic and 0.5kPa PDMS +/- addition of DAPT after immunofluorescence staining are displayed on the right panel (VE-cadherin in green, scale bar 50μm).

### Decreased substrate stiffness elevates NECD trans-endocytosis but not general endocytosis

To investigate the role of Notch receptor-ligand binding in increased Notch signaling activity on softer substrates, we performed a trans-endocytosis assay [28]. Separate cell populations were transfected with a Notch1-citrine fusion plasmid, or a doxycycline controlled Dll4-mCherry fusion plasmid. Dll4 expression was induced by adding doxycycline to the co-culture of both transfected cells. Analysis was performed in the areas where the signals of Notch1 receptor and Dll4 ligand overlap at the cell-cell contacts of a signal-sending and a signal-receiving cell using ImageJ. To quantify trans-endocytosis and examine the stiffness effects, the colocalization of Notch receptor and ligand was analyzed both in the form of intensity analysis of the overlay areas, and as a correlation analysis using Pearson’s r coefficient. Trans-endocytosis peaked after 6h of incubation as described by Shaya et al [28]. Results show that trans-endocytosis increases on soft substrates, which is reflected in an increased intensity of the interaction area, as well as in a higher colocalization coefficient, the Pearson’s r value (Figure 6A). A transferrin endocytosis assay was performed on the substrates to exclude a stiffness effect on endocytosis in general. The results show that softer substrates do not enhance general endocytosis, so the increased trans-endocytosis cannot be attributed to general increase in endocytosis. Thus, trans-endocytosis seems to selectively exhibit mechanosensitivity in cell-cell contact-dependent receptor binding. The same experiments were performed with the endothelial cell line MCEC-WT for comparison, showing similar results and the same effect; the softer the substrate, the more trans-endocytosis occurs upon cell-cell contact (Supplementary Figure S3).

**Figure 6.**
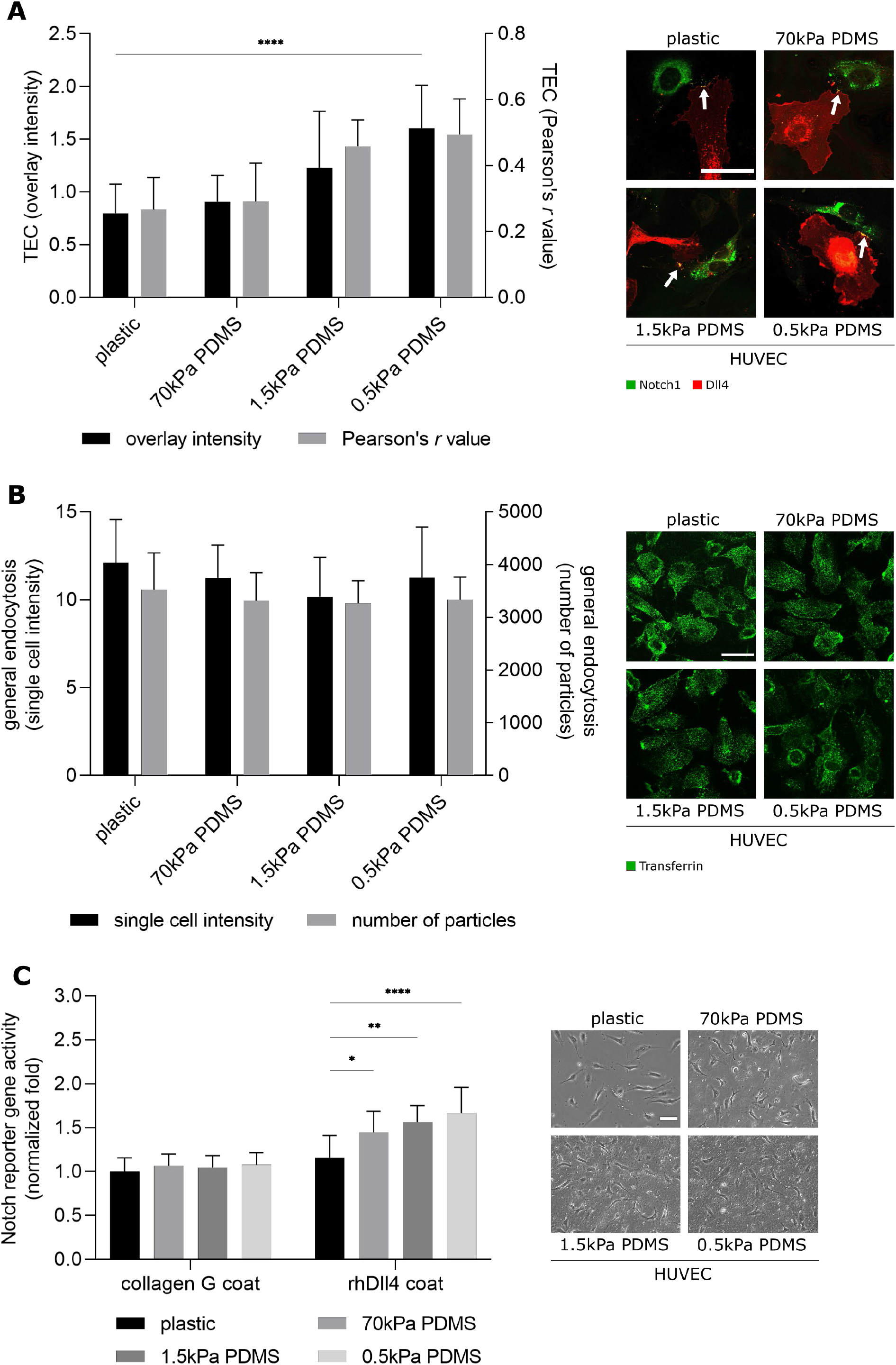
Trans-endocytosis is increased on softer substrates and cell-cell contacts are dispensable for Notch activation with rhDll4. (A) Overlay intensity and Pearson’s r value in areas of Notch receptor ligand interactions in the course of trans-endocytosis. Overlay areas are indicated by the white arrows. HUVEC cells were transfected separately with a citrine-coupled Notch1 plasmid and a mCherry-coupled Dll4 plasmid. Trans-endocytosis was quantified at cell-cell contacts in ≥30 cells per substrate condition in three independent experiments. Data is presented as a bar plot on the left panel (mean value ± SEM, two-way ANOVA with Sidak’s multiple comparison test, ****P<0.0001). Representative images of TEC are shown on the right panel (Notch1 expressing Notch receiver cells are shown in green, Dll4 expressing Notch sender cells are shown in red, scale bar 50μm). (B) General endocytosis in HUCEC. Cells were seeded on substrates with different stiffness and a transferrin endocytosis assay was conducted. Intensity and number of particles in individual cells is presented in a bar plot as means ± SEM of ?300 cells per substrate condition (Sidak’s corrected two-way ANOVA, *P<0.1). (C) Normalized reporter gene activity in non-confluent HUVEC cells with and without Notch activation via a rhDll4 coat, determined by detection of cellular luciferase levels under control of the TP1-luc Notch reporter. Bar plots were generated by evaluation of reporter gene assays on substrates with different stiffness shown on the left panel (one-way ANOVA followed by Tukey’s multiple comparison test, *P<0.1, **P<0.01, ****P<0.0001). Representative images of the cell density are shown on the right panel (scale bar 50μm).

Nevertheless, after activation with the rhDll4 coat, Notch recipient cells also showed increased Notch signaling activity even without trans-endocytosis taking place. Therefore, to investigate whether the stiffness effect of the Notch signaling pathway is also detectable independently of cell-cell contacts, and thus independently of trans-endocytosis only by Notch activation with rhDll4, a reporter gene assay was performed in nonconfluent HUVECs, instead of confluent cells as in Figure 1A. We observe a significant increase in reporter gene activity after Notch activation by rhDll4 on softer substrates (Figure 6C), where the rhDll4 coat is evenly distributed on all substrates (Figure S1). In contrast, no change in reporter gene activity was evaluated on the substrates without Notch activation (collagen G coat). Thus, cell-cell contacts contribute to Notch activity but are not necessary for a Notch signaling effect in this setting. Although both activation methods of the Notch signaling pathway showed increased Notch signaling activity on softer substrates, little is known about cell contact-independent Notch activation. Thus, activation by rhDll4 and the subsequent pathway may be affected by stiffness changes differently than activation by a neighboring cell.

## Discussion

The Notch signaling pathway plays multiple and crucial roles in developmental and pathological processes [29–31]. Many of these scenarios are related to changes in the composition and the biomechanical features of the extracellular matrix. During angiogenesis the Notch signaling pathway regulates cell fate decisions such as migration, proliferation, and differentiation, essential for vascular development and angiogenesis [10, 32]. Endothelial Notch signaling can be activated by VEGF, and, at the same time, VEGF controls matrix composition, causing local ECM softening or stiffening [12, 33]. Previous studies on biophysical aspects of Notch signaling focus on a pulling force exerted by the bound receptor [34] that is a prerequisite for Notch receptor cleavage. To date, however, little is known on whether the Notch signaling pathway is modulated by substrate stiffness. To address this question, we used an endothelial cell model based on synthetic PDMS substrates with tunable stiffness Interestingly, nuclear NICD localization and transcriptional activity of Notch both increase with decreasing substrate stiffness. This phenomenon occurs irrespective of whether Notch activation was achieved by surface bound ligand (rhDll4) or Dll4 overexpressing sender cells (Fig. 1).

Endothelial cells sense changes of matrix properties via cell-matrix adhesion proteins, such as integrin β1 [16, 33], which also plays a crucial role in regulating cellular adhesion to enable migration and proliferation [23, 33]. Since integrins have been previously located both up- [35] and downstream of Notch in signaling pathways [7], we first investigated this issue. For integrin β1, it has already been shown that the overall levels do not change with different substrate stiffnesses, but that the active state of integrin β1 increases on softer substrates [22]. We confirm these observations in our model. Furthermore, our data show that cell-matrix adhesion is directly influenced by the Notch signaling pathway. Notch inhibition by DAPT treatment significantly reduced the integrin β1 intensity of the activated form, although substrate-dependent activation was further observed. (Fig. 2). These results corroborate the findings of Hodkinson et al, which show that Notch1, through NICD cleavage and subsequent R-ras binding, can induce a conformational change in the surface-bound integrin from a low affinity state to a high affinity state, mobilizing the active form [7]. This clearly positions integrins downstream of Notch. Despite Notch inhibition, a continuous increase in integrin β1 levels on softer substrates can be observed, however, on a much lower level (Fig. 2). Thus, either pharmacological Notch inhibition was not complete, or integrin activity is additionally influenced by substrate stiffness via an alternative pathway.

The role of the Yes-associated protein YAP as a mechano-transducer and the related mechanosensitivity of the YAP/TAZ signaling pathway are well known and studied. Upon YAP activation, for example by higher ECM rigidly or substrate stiffness, YAP/TAZ are not phosphorylated and can thus be relocated from the cytoplasm to the nucleus, where binding to a transcription factor regulates genes necessary for cell migration and proliferation [24, 36]. Accordingly, we investigated, how the YAP/TAZ pathway relates to stiffness modulated Notch signaling. As expected, YAP activity decreased with lower stiffness of the substrate (Fig.3). It has been previously shown that activation of YAP reduces Notch signaling by repression of the ligand Dll4, while knockdown of YAP increases Dll4 [37]. However, this mechanism is not sufficient to explain the correlation between substrate stiffness and Notch activity in our model, since we examined Notch activation after co-culture with a stably overexpressing cell line. Additionally, increased activity of Notch with lower stiffness was also observed after activation by rhDll4, which is independent of YAP activity. There also seems to be no feedback loop between Notch and YAP translocation, since DAPT treatment did not alter stiffness dependence of YAP translocation to the nucleus (Fig. 3). However, we cannot exclude concomitant activation of YAP and Notch on a transcriptional level, which has been described previously [2].

In addition to the cell-matrix adhesions, a stiffness effect has already been established for cell-cell contacts. On stiffer substrates, VE-cadherin junctions are wider and discontinuous, whereas on soft substrates they are narrower but continuous [38]. We corroborate this finding by analyzing VE-cadherin patterns through mean branch length evaluation. The VE-cadherin junctions on plastic show a branched, comb-like structure with larger branch length in contrast to the narrower junctions on softer substrates with smaller branch length, which appear more distinct and continuous (Fig. 4). A role for cadherins in the regulation of the Notch signaling pathway was previously described by Kwak et al, who demonstrated that cadherin-based adherens junctions control Notch-γ-secretase interactions [39]. In this context, the changes in VE-cadherin junctions on the soft substrates could be related to the increased Notch activities. However, it should be noted that the VE-cadherin patterns on the changing substrate stiffnesses do not affect the intensity level or VE-cadherin trafficking (Fig. 4). Furthermore, Notch signaling is regulated by the cell-cell contact area [2, 28], as signaling is directly linked to the contact area of two interacting cells [28]. By destabilizing and blocking the VE-cadherin cell-cell contacts, we show that, as expected, Notch activity is significantly reduced. This effect is, however, independent of substrate stiffness (Fig. 5). With these results, it should be noted that the Notch receptor has a Ca^2+^ dependency, which means that the use of EGTA to destabilize the cell-cell contacts could also destabilize the receptor, leading to Notch cleavage and activation [26, 27]. However, analysis of NICD levels after EGTA treatment shows that in our model a reduced NICD nuclear intensity was detected and thus no EGTA mediated Notch activation occurred, additionally noting that EGTA was dissolved in medium here in contrast to other studies [26]. Blocking the Notch signaling pathway leads to a uniform decline of VE-cadherin on all substrates. The change of junctional patterns due to variation of stiffness is not influenced by DAPT treatment (Fig. 5). These findings argue against an upstream role of VE-Cadherin for stiffness regulation of Notch signaling.

By endocytosis of the Notch ligand after binding to the receptor into the signal-sending cell, the accumulation of the Notch receptor on the cell surface of the signal-receiving cell can be counteracted on the one hand and serves primarily for receptor activation on the other[11, 40]. During trans-endocytosis, the extracellular part of the Notch receptor bound to the Notch ligand is pulled into the ligand-presenting cell, thus releasing the intracellular part of the receptor (activated form), which can be released from the membrane in a further cleavage event [6]. Our results from the TEC assay on the different substrates show that trans-endocytosis is significantly enhanced on soft substrates which is consistent with the enhanced Notch activity on softer substrates. A stiffness effect on the general endocytosis can be excluded. Taken together, the observations demonstrate a mechanosensitivity of the Notch signaling pathway that may in part be due to trans-endocytosis. Interestingly, activation of the Notch signaling pathway in a cell-cell contact independent manner by using rhDll4 as a stimulus for subconfluent single cells, we can also show that the cell-cell contacts are to a certain degree dispensable in this setting. Upon Notch pathway activation by a Dll4 coat, the stiffness dependence of Notch can still be shown, whereas a different activation mechanism takes effect and thus the Notch receptor activation and the influence of substrate stiffness on the Notch signaling pathway may be different than after activation by cell-cell contact. While the Notch pathway after activation by interaction with a ligand presenting cell shows a mechanical aspect of the pulling movement of the ligand into the sender cells, the mechanism of mechanosensitivity of the Notch pathway after rhDll4 activation and thus without trans-endocytosis and pulling movement is unknown and possibly subject to a different mechanical force.

These findings might be of importance to better understand the effect of matrix remodeling on endothelial cell behavior and adjustment of their gene expression [12, 17]. Matrix remodeling increases the elasticity of the matrix and thus decreases the stiffness of the ECM [17, 18], thus triggering a feedback loop. With Notch being one of the main regulators of endothelial cell fate decisions as well as being the key mediator of angiogenesis by controlling tip and stalk cell differentiation [11, 41], our results provide further insights on the consequences of normal and pathological matrix changes on endothelial behavior during angiogenic sprouting in development and maintenance.

## Supporting information

Supplements - Matrix stiffness regulates Notch signaling activity in endothelial cells

## Acknowledgements

This project was funded by the Deutsche Forschungsgemeinschaft (DFG, German Research Foundation) - Project-ID 201269156 - SFB 1032 (Project B8). DS acknowledges the support of the European Research Council (ERC) under the European Union’s Horizon 2020 research and innovation programme (Grant agreement No. 682161).

