## Supplements - Matrix stiffness regulates Notch signaling activity in endothelial cells for "Matrix stiffness regulates Notch signaling activity in endothelial cells"

### Supplementary figures

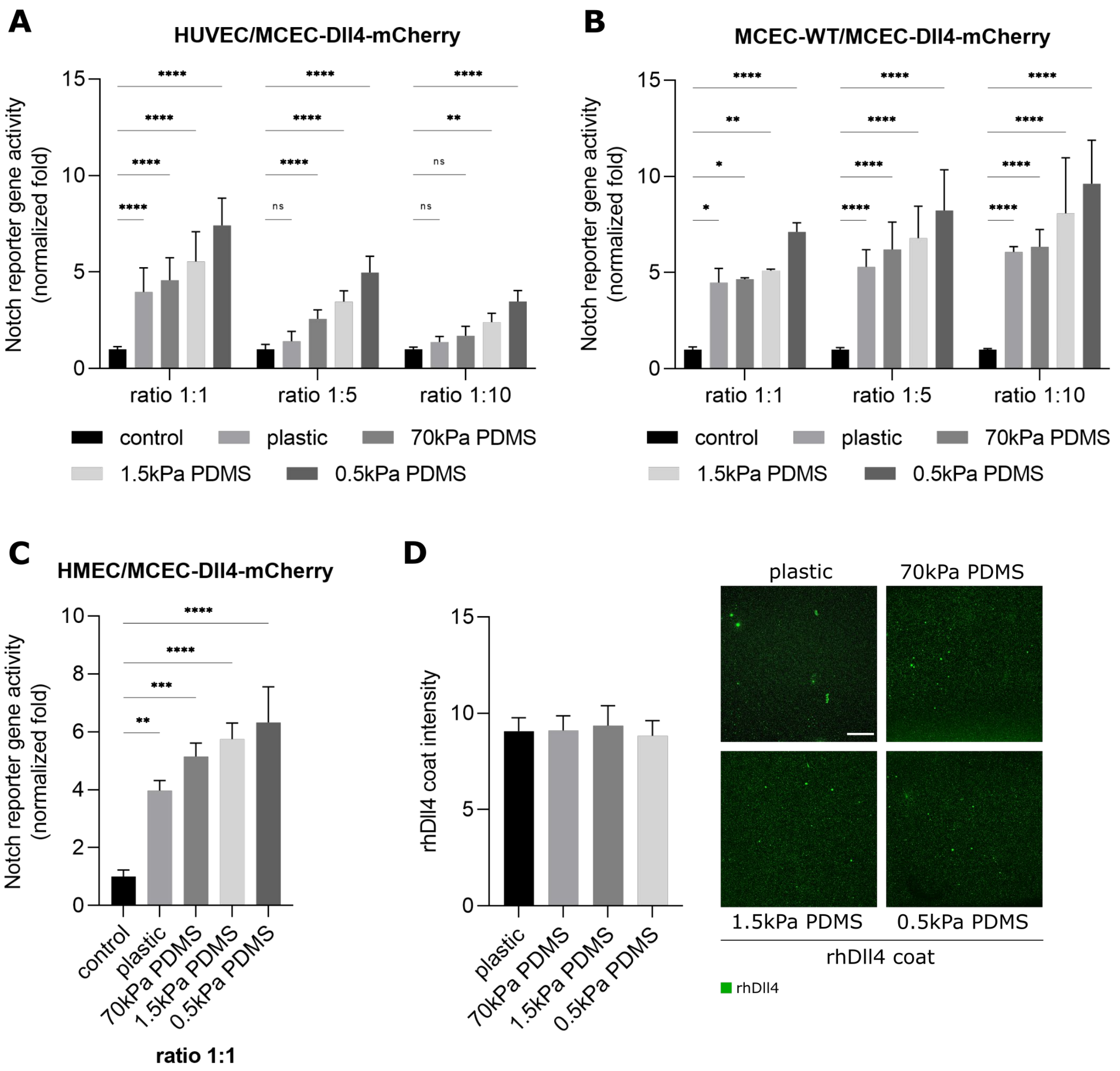


**Figure S1. Notch activation by co-culture of Notch sender and receiver cells increases in soft substrates in all Notch receiver cells but is dependent on the seeding ratio.**

(A,B) Normalized fold Notch activity in endothelial co-cultures of HUVEC/MCEC-Dll4-mCherry cells (A) and MCEC-WT/MCEC-Dll4-mCherry cells (B) in seeding ratios of 1:1, 1:5 and 1:10. Bar plots were generated by evaluation of reporter gene assays on substrates with different stiffnesses (two-way ANOVA followed by Tukey's multiple comparison test, **P<0.01, ****P<0.0001). (C) Normalized fold Notch activity in HMEC/MCEC-Dll4-mCherry co-culture ratio 1:1, outlined in a bar plot (mean ± SEM, Tukey's corrected one-way ANOVA, **P<0.01, ***P<0.001, ****P<0.0001). (D) PDMS substrates were coated with rhDll4 and stained for Dll4 (shown in green). rhDll4 binding efficiency was compared by evaluation of intensity and the number of particles, summarized in a bar plot (mean ± SEM, two-way ANOVA followed by Tukey's multiple comparison test).

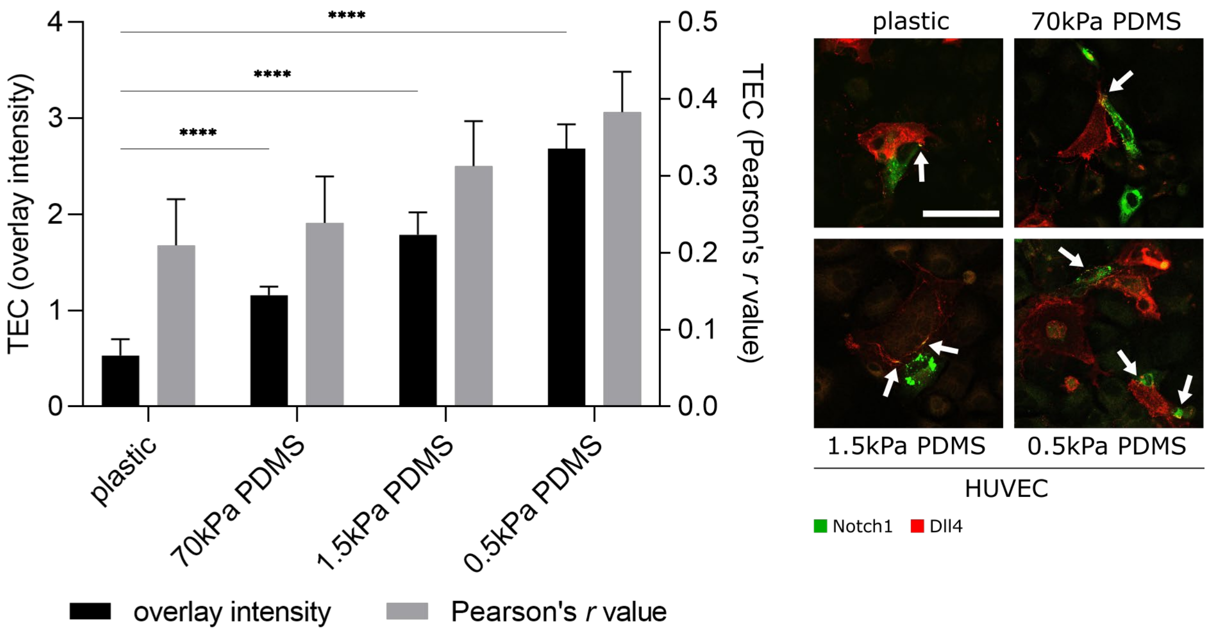


**Figure S2. Trans-endocytosis is also in MCEC-WT cells increased on softer substrates.**
Overlay intensity and Pearson's r value in areas of Notch receptor ligand interactions in the course of trans-endocytosis. Overlay areas are indicated by the white arrows. MCEC-WT cells were transfected separately with a citrine-coupled Notch1 plasmid and a mCherry-coupled Dll4 plasmid. Notch1 expressing Notch receiver cells are shown in green, Dll4 expressing Notch sender cells are shown in red. Trans-endocytosis was quantified at cell-cell contacts in ≥30 cells per substrate condition in three independent experiments. Data is presented as a bar plot (mean value ± SEM, two-way ANOVA with Sidak's multiple comparison test, ****P<0.0001).

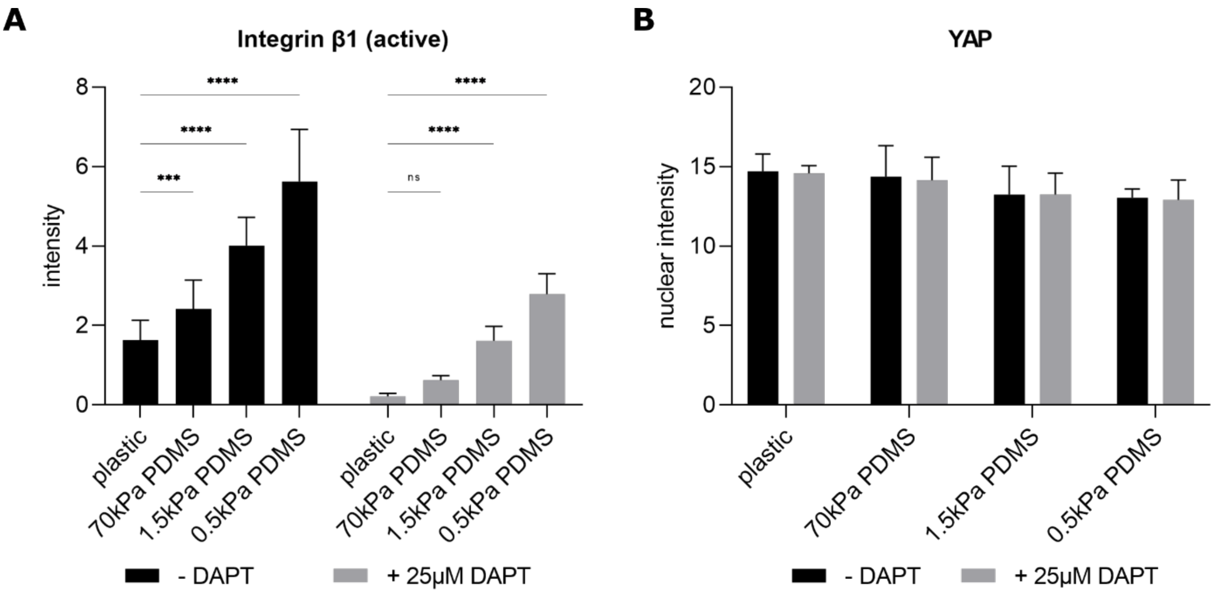


**Figure S3. Integrin β1 intensity in MCEC-WT cells is dependent on substrate stiffness as well as the Notch signaling pathway, whereas the nuclear intensity of YAP does not change after Notch inhibition by DAPT.**
(A,B) MCEC-WT cells were seeded on varying substate stiffness, treated with 25µM DAPT for 24h and stained for either the activated form of integrin β1 or YAP. The mean overall intensity for integrin β1 and the nuclear intensities for YAP ± SEM of untreated and treated cells are summarized in bar graphs (Tukey's/Sidak's corrected two-way ANOVA, ***P<0.001, ****P<0.0001).
